# Canvas SPW: calling *de novo* copy number variants in pedigrees

**DOI:** 10.1101/121939

**Authors:** Sergii Ivakhno, Eric Roller, Camilla Colombo, Philip Tedder, Anthony J. Cox

## Abstract

**Motivation:** Whole genome sequencing is becoming a diagnostics of choice for the identification of rare inherited and *de novo* copy number variants in families with various pediatric and late-onset genetic diseases. However, joint variant calling in pedigrees is hampered by the complexity of consensus breakpoint alignment across samples within an arbitrary pedigree structure.

**Results:** We have developed a new tool, Canvas SPW, for the identification of inherited and *de novo* copy number variants from pedigree sequencing data. Canvas SPW supports a number of family structures and provides a wide range of scoring and filtering options to automate and streamline identification of *de novo* variants.

**Availability:** Canvas SPW is available for download from https://github.com/Illumina/canvas.

**Supplementary information:** Supplementary data are available at *Bioinformatics* online.

## 1 INTRODUCTION

The advent of affordable high-throughput sequencing enables the characterization of genes implicated in a wide range of genetic diseases and makes it possible to provide diagnosis in clinical settings as exemplified by Genomics England 100k Genomes Project for the National Health Service (https://www.genomicsengland.co.uk). Whole genome sequencing of families is becoming a standard approach for identifying highly penetrant variants that cause rare disease, such as *de novo* or recessive mutations. Accurate variant and genotype calling is crucial for successful identification of such disease-causing mutations (Acuna-Hidalgo et al. 2016). Unfortunately, false positive and negative results can occur due to technical artifacts or reduced sequencing coverage, which especially impact copy number variants (CNVs) identified through read depth estimation (Teo et al. 2012). CNV calling accuracy in families can be improved over single-sample calling by incorporating pedigree structure into the genotyping model to ensure that copy number genotypes are consistent with Mendelian inheritance and low rates of *de novo* mutation. While a number of tools have been developed for the identification of germline CNVs from sequencing data (Boeva et al., 2011, Abyzov et al., 2011 and Liu et al., 2016), most of them are limited to variant calling in single samples. Even those designed for family-based CNV detection are restricted to only deal with parent-offspring trios (Liu et al., 2016). To expand the number of analyzable family structures and provide explicit and easy-to-interpret *de novo* variant calls, we have developed a new workflow, Canvas SPW (Small Pedigree Workflow), for germline and *de novo* variant calling in pedigrees. In addition to trios, Canvas SPW processes quads and can also perform joint variant calling in medium-size sample batches.

## 2 METHOD

### Outline

Canvas SPW comprises five distinct modules designed to (1) process aligned read data and estimate depth in coverage bins, (2) perform outlier removal and normalization of depth estimates, (3) partition bins into segments of uniform copy number, (4) calculate associated allele counts from single nucleotide variants (SNVs) and (5) assign germline and *de novo* copy numbers. While initial multi-sample data processing and normalization steps are extensions of single-sample methods described elsewhere (Roller et al., 2016), segmentation and variant calling steps have been specifically developed for multi-sample pedigree-structured inputs. We briefly describe them below. More information is available in the Supplementary Material, Section 1.

### Segmentation

The input to the segmentation module is a data matrix produced by processing the aligned read data from all samples. Each row of the input data matrix represents a single genomic bin with the same genomic coordinates across all samples. In turn, each column represents the normalized depth estimate across all genomic bins for a given sample. A Hidden Markov Model (HMM) with multivariate negative binomial emission distribution uses this matrix for partitioning. HMM hidden states are initialized to approximately follow copy number states (exact CN assignment is done at the variant calling stage). First, the Expectation Maximization algorithm is used to optimize parameters of the emission and transition distributions. Next, the Viterbi algorithm is used to derive the final partitions.

### Variant calling and output

For each segment determined by the HMM segmentation module, Canvas SPW uses the distribution of coverage across the various bins along with the allele-specific depths at each SNV within the segment to assign a copy number. As a rule, allele-specific depths are only used when a segment contains enough SNV loci; allele-specific depths are not use in small segments containing only a handful of SNV sites. A probabilistic model is fitted to estimate likelihood (L) of copy number (CN) assignments within a pedigree given coverage data (D) *L*(*CN_m_*, *CN_f_*, *CN_c_/D*)~ *P(D_m_*/*CN_m_*) *P(D_f_*/*CN_f_*) *P(D_c_*/*CN_c_*) × *P*(*CN_c_*/*CN_m_*, *CN_f_*), where the last term incorporates both the Mendelian transmission probabilities and the estimated *de novo* rate of CNVs. Likelihood evaluation is done using exhaustive enumeration of all possible *CN* assignments within the pedigree up to the maximal CN threshold. A joint probability table from the model with the maximum likelihood is used to estimate single sample and *de novo* quality scores for variant calls and every variant inconsistent with Mendelian inheritance is assigned a *de novo* flag. A VCF 4.1 compliant file with common and *de novo* CNV calls is produced at the end of each Canvas SPW run.

### Implementation and performance

Canvas SPW is implemented in the C# programming language and can be run on Linux systems using mono/.NET Core or on Windows systems under the .NET. Using a Linux system with 32 cores and peak RAM consumption of less than 10G, the Canvas SPW runtime on a trio and a quad pedigree with 40X sequencing coverage per-sample was 1.3h and 2.1h respectively.

## 3 RESULTS

Two key performance characteristics of Canvas SPW were evaluated: (1) ability to accurately call inherited germline variants and (2) correct *de novo* variant detection. To accomplish this we have created a number of pedigree sequencing samples using Platinum Genomes (PG) dataset (Eberle et al., 2016) that include: (1) normal trios, (2) negative control replicates of a single sample, (3) *de novo* enriched trio and quad where parents are derived from the same sample and (4) pedigree simulation through haplotype down-sampling. The latter is an adaptation of the previously described tHapMix simulation framework for somatic variants (Ivakhno et al., 2016) to germline CNVs within a pedigree relationship structure. Truth sets were generated by merging structural variants found in PG data using orthogonal variant calling tools. Further validation of these truth sets was done by checking for full consistency with Mendelian inheritance and comparison with TruSeq synthetic long read data (full details of the assessment methodology and results are available in Supplementary Material, Section 2). Estimation of CNV calling accuracy, precision and recall was performed as previously reported using genome-wide base-pair CN concordance (Roller et al., 2016). We assessed the performance of Canvas SPW against a range of existing CNV calling tools: Control-FREEC (Boeva et al., 2011), CNVnator (Abyzov et al., 2011) and TrioCNV (Liu et al., 2016). All tools were tested for germline CNV calling accuracy, precision and recall. TrioCNV was also evaluated for its ability to call *de novo* variants. The assessment was done separately on four real synthetic (Table 1) and six haplotype-simulated pedigrees (Table 2).

**Table 1.**
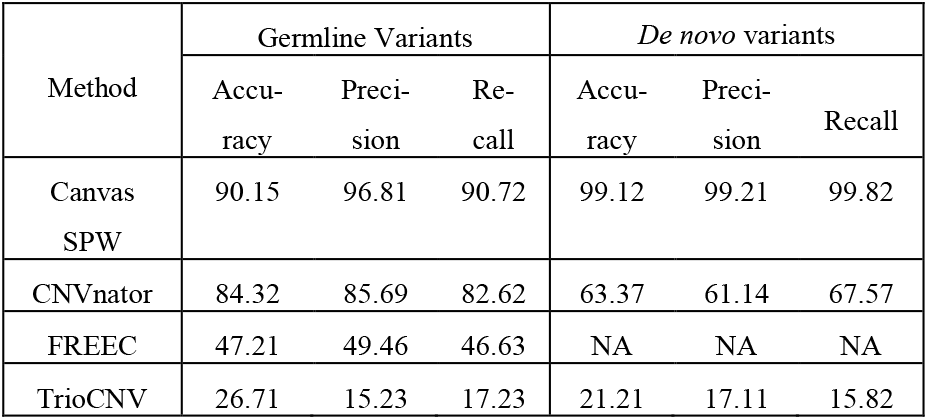
CNV calling performance metrics for real synthetic pedigrees

**Table 2.** CNV calling performance metrics for haplotype simulated pedigrees

| Method | Germline Variants |  |  | De novo variants |  |  |
| --- | --- | --- | --- | --- | --- | --- |
|  | Accu-racy | Preci-sion | Re-call | Accu-racy | Preci-sion | Recall |
| Canvas SPW | 91.21 | 94.39 | 88.76 | 98.25 | 98.89 | 97.14 |
| CNVnator | 75.24 | 87.45 | 77.82 | 62.19 | 49.14 | 64.57 |
| FREEC | 40.82 | 43.29 | 42.38 | NA | NA | NA |
| TrioCNV | 22.34 | 12.86 | 13.93 | 24.57 | 15.27 | 13.49 |

Canvas SPW showed superior performance in comparison with existing tools for both inherited and *de novo* germline variants. It also outperforms the single-sample germline Canvas workflow (Supplementary Material, Table 2), suggesting that joint CNV calling with pedigree information not only enables *de novo* variants detection, but also improves performance on inherited germline variants. This is particularly true for the recall where the pooling of sequencing coverage from multiple samples amplifies signal, thereby decreasing the number of false negative calls. To conclude, Canvas SPW provides fast, accurate and easy to use workflow for the identification of inherited and *de novo* germline CNV variants in pedigrees.

## Acknowledgements

We thank Stephen Tanner and Lisa Murray for valuable discussions.

